# Akaby - cell-free protein expression system for linear templates

**DOI:** 10.1101/2021.11.03.467179

**Authors:** Wakana Sato, Judee Sharon, Christopher Deich, Nathaniel Gaut, Brock Cash, Aaron E Engelhart, Katarzyna P Adamala

## Abstract

Cell-free protein expression is increasingly becoming popular for biotechnology, biomedical and research applications. Among cell-free systems, the most popular one is based on *Escherichia coli* (*E. coli*). Endogenous nucleases in *E. coli* cell-free transcription-translation (TXTL) degrade the free ends of DNA, resulting in inefficient protein expression from linear DNA templates. RecBCD is a nuclease complex that plays a major role in nuclease activity in *E. coli*, with the RecB subunit possessing the actual nuclease activity. We created a *RecB* knockout of an *E. coli* strain optimized for cell-free expression. We named this new strain Akaby. We demonstrated that Akaby TXTL successfully reduced linear DNA degradations, rescuing the protein expression efficiency from the linear DNA templates. The practicality of Akaby for TXTL is an efficient, simple alternative for linear template expression in cell-free reactions. We also use this work as a model protocol for modifying the TXTL source *E. coli* strain, enabling the creation of TXTL systems with other custom modifications.

## Introduction

Cell-free transcription-translation (TXTL) has been gathering increased attention in research and industry, due to its versatile application potential in synthetic biology. Those applications include, gene circuit testing[1], artificial cell systems[2–4], protein evolution[5], and enzymatic reaction optimizations[6,7]. The combination of bacteriophage RNA polymerase T7 and *Escherichia coli* (*E. coli*) crude extract is the most popular TXTL system[8]. In TXTL, the proteins are expressed from gene coding DNA in either circular plasmids, or linear DNA fragments. There have been tremendous characterizations and optimizations to improve nearly all aspects of TXTL[1,9,10].

Translation of linear DNA templates is of great importance to both research and practical applications. Eliminating the need to clone plasmid DNA, TXTL can be used to significantly increase the speed of the design – build – test cycle in bioengineering. Since the bacterial TXTL degrades linear DNA templates, two nuclease inhibitors are widely used to overcome this problem: GamS or Chi6 (short DNA containing six Chi sites). Neither of these inhibits the nuclease activity completely.

RecBCD is a nuclease complex endogenous to *E*.*coli. RecB* contains the nuclease and the helicase domains; *RecC* is implicated to be responsible for Chi sequence recognition; and *RecD* contains the helicase domain[11].

RecBCD plays a vital role in maintaining endogenous genome quality and integrity. One of the most common DNA damage mechanisms is a DNA double-strand break caused by a variety of reasons, such as ionizing radiation or DNA replication errors. To repair this damage, RecBCD binds blunt-ended DNA termini and converts the template into a duplex DNA possessing a 3’-terminated ssDNA tail to initiate homologous recombination. The critical component in this process is the octameric regulatory DNA sequence called Chi (5’-GCTGGTGG-3’)[12,13]. RecBCD terminates the conversion of dsDNA into ssDNA at the Chi sequence. Since TXTL consists of *E. coli* crude extract, endogenous RecBCD remains active in the TXTL reactions. It has been previously demonstrated that the supplementation of Chi6 DNA enhances DNA stability and protein expression in TXTL[14]. This finding strongly suggests the association of RecBCD in *E. coli* crude extract with DNA stability and gene expression in TXTL.

Along with the Chi sequence-based DNA degradation, RecBCD also possesses random nuclease activity for blunt-ended DNA fragments. That random nuclease activity initially evolved to defend their own genome from invading DNA, such as lambda and T4 bacteriophages[11]. GamS, a bacteriophage lambda encoded protein, evolved as a counter-strategy of RecBCD to protect phage DNA by forming a stable complex with RecBCD[15–17] GamS forms a double-stranded DNA mimetic and works as a competitive inhibitor for RecBCD[18]. To protect free-end linear DNA templates in TXTL, GamS has been used as a supplementation of TXTL to increase the protein expression from the gene template of linear PCR fragments[1]. Even though GamS has been used to protect linear DNA fragments in TXTL, it is still unclear which nuclease in TXTL plays the vital role in reducing protein expression from linear templates.

Both GamS and Chi6 DNA protection protocols require adding additional components to TXTL. This not only adds extra steps, costs and failure points to the reaction, but it also uses some of the limited component volume. TXTL reactions are typically set up with very little volume left for additives and templates. We provide a solution to those problems by engineering an *E. coli* strain optimized for cell-free TXTL production with inactivated RecBCD nucleases.

It has been previously demonstrated that knocking-out or -down RecBCD nuclease in *E. coli* strains result in cell-free lysate that shows improved template stability[19,20]. We decided to build on that work, using the DH5α *E. coli* strain optimized for TXTL, and creating a strain that will be freely available to the whole community. Using state of the art mutagenesis techniques provides an optimized protocol for future engineering of custom TXTL strains with specific properties.

We engineered the *RecB* knockout *E. coli* strain, which we call Akaby. The name was inspired by the Arcade game Pac-Man, where Akabei is the Japanese name for the Blinky ghost nemesis of Pac-Man. Pac-Man figure is commonly used in simplified schemes to indicate nucleases. The Akabei ghost incapacitates Pac-Man, analogous to the Akaby strain with an incapacitated major nuclease.

We demonstrated that RecB is the strongest nuclease affecting unprotected linear DNA fragment stability in TXTL, and Akaby provides protection from linear DNA degradation. Using Akaby cell-free extract for TXTL can be a simple, efficient choice for linear DNA-friendly TXTL platforms.

## Results and Discussion

### *RecB* disruption in *E. coli* DH5α

RecB is the RecBCD subunit that possesses nuclease activity. We deleted the *RecB* gene from the genome of *E. coli* DH5α by using the λ red system described previously, with minor modifications[21]. The kanamycin-resistant gene (Km^R^) was chosen as a selection marker. On the *E. coli* DH5α genome, *RecB* is located between *ptrA* and *RecD* (**Fig. 1A**). Therefore, we designed primers for Km^R^ PCR amplification with 120 nt homology extensions, which are complementary with the upstream or downstream of *RecB* in *E. coli*. These homology extensions promote the replacement of *RecB* with Km^R^ on the *E. coli* genome.

**Fig.1.**
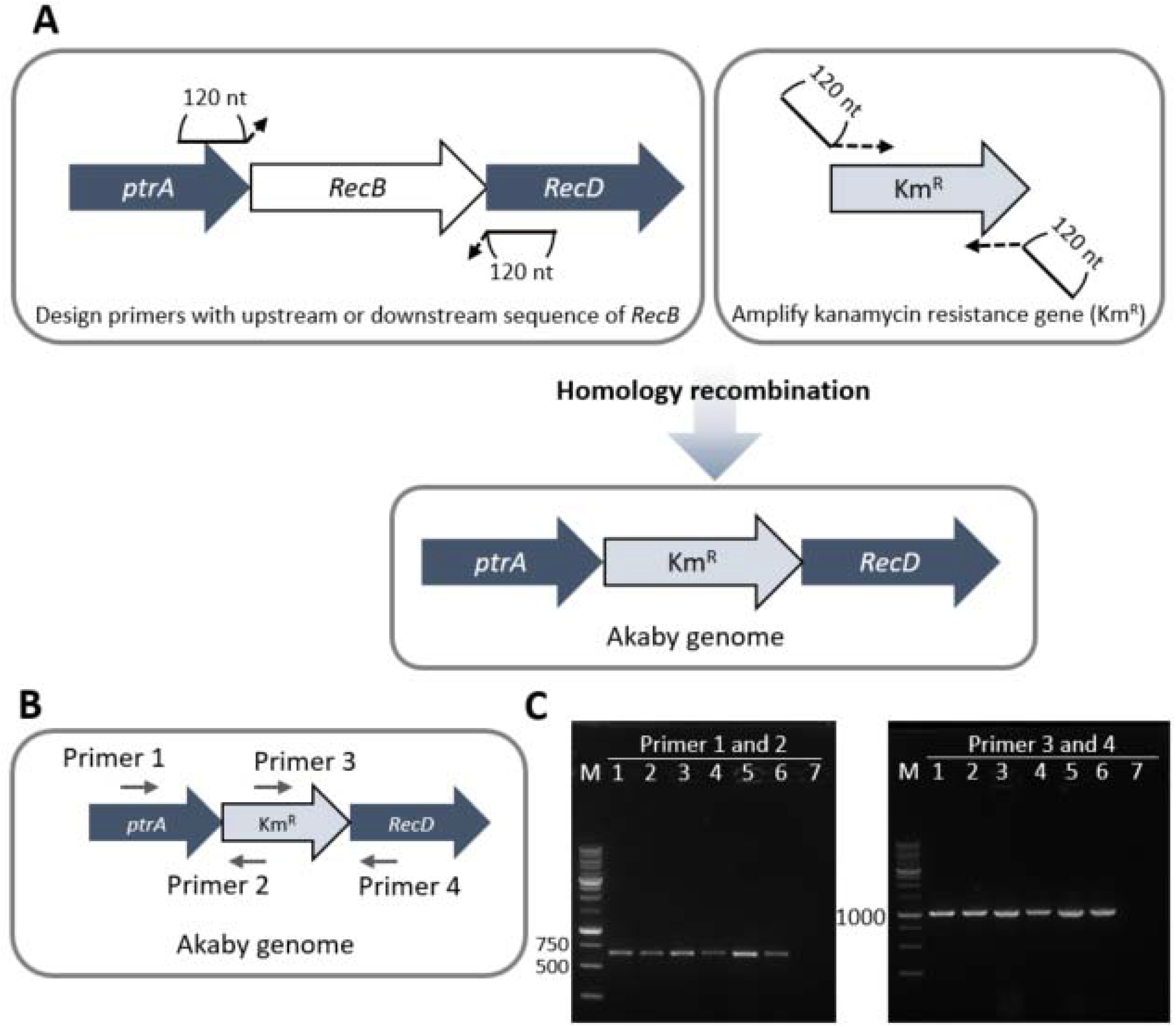
Overview of the *RecB* knockout experiment. (A) The primer designs for Kanamycin resistant gene (Km^R^) PCR. The primers contains 120nt upstream or downstream sequences of *RecB* in DH5α genome. (B)The primer designs for colony PCR. Primer 1 and 4 have complement sequences outside of *RecB* in DH5α genome. Primer 2 and 3 have complement sequences within Km^R^. (C) Agarose gel images of colony PCR. 1% agarose gel was run at 125 V for 45min, stained with SYBR safe DNA Gel stain. Expected product size: 661 bp (left gel) and 1035 bp (right gel). Lane M: 1 kb DNA ladder (Goldbio, D010-500) lane 1∼6: *RecB* replaced DH5α colonies, 7: wild type DH5α colony.

After the recombineering procedure, the successful mutant was verified through colony PCR. For the colony PCR, the insert-(Primer 2 and 3) and locus-(Primer 1 and 4) specific primer pairs were used (**Fig. 1B**). While no PCR product was detected for the untreated *E. coli* colony (**Fig. 1C**, lane 7), all six analyzed mutant colonies produced the expected size of PCR products. We thus confirmed that the successful mutant strain that contains the genome replaced *RecB* with Km^R^. We prepared Akaby cell extract from a glycerol stock generated from a single colony among the six confirmed mutants for later experiments.

### Akaby TXTL increased eGFP fluorescence from linear templates

eGFP expressions with circular and linear templates were compared to evaluate the property of Akaby crude extract in cell-free transcription-translation (TXTL) (**Fig. 2**). For linear templates, we used eGFP plasmids digested with BamHI. In that plasmid, BamHI cuts at position 182 nt after the eGFP coding sequence, thus linearizing the eGFP plasmid without affecting the open reading frame. To further study the robustness of the system, we used eGFP templates under two different T7 RNA polymerase promoters, the canonical T7 promoter and the new high yield T7Max promoter[22]. Because TXTL components generate a low level of endogenous fluorescence in the green channel, we used a no template control (NTC) as the background fluorescence benchmark. As a comparison to the Akaby cell extract, we used the Rosetta 2 cell extract that is most commonly used for the TXTL experiments[23].

**Fig. 2.**
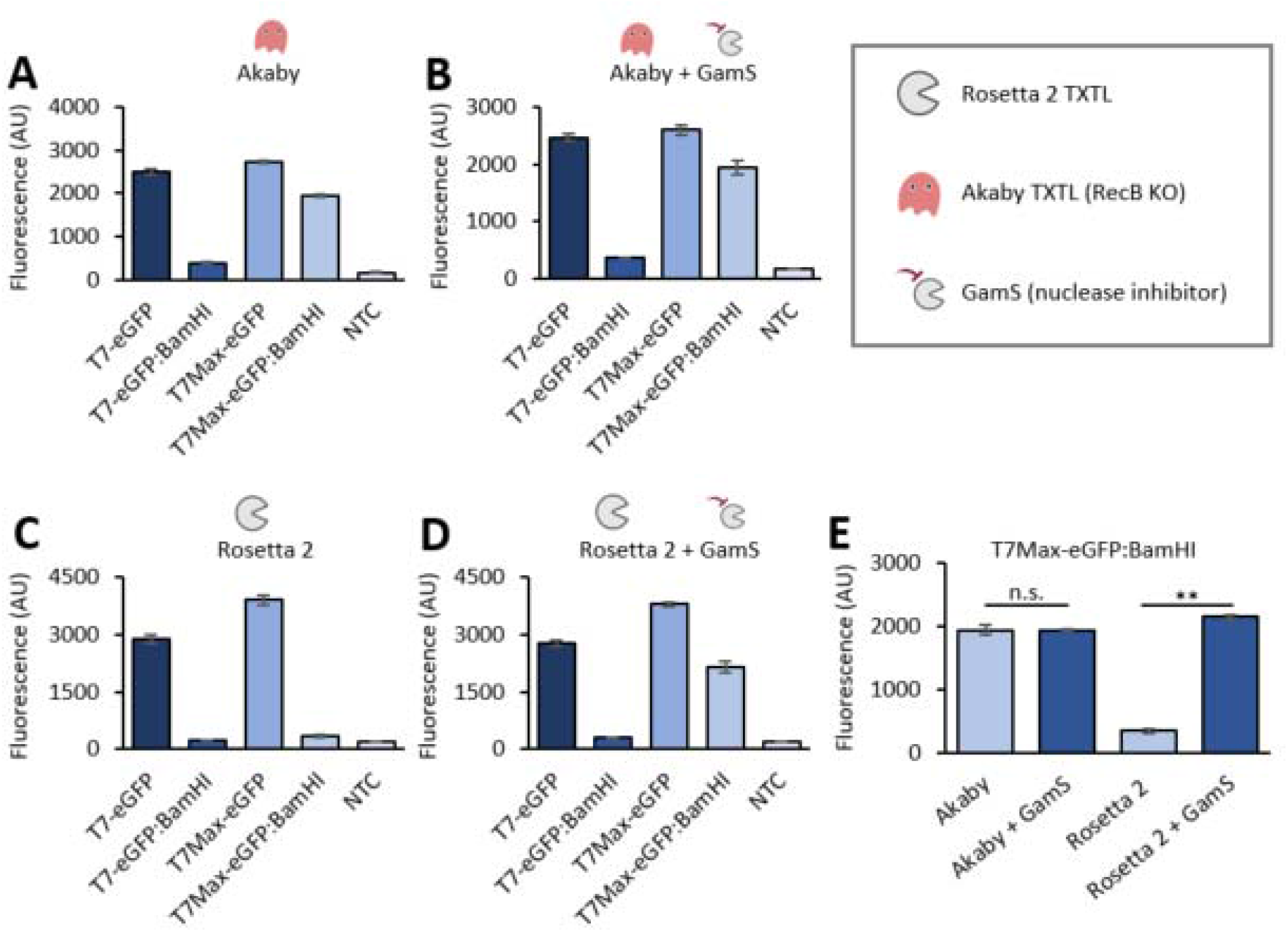
eGFP expression comparison between Akaby and Rosetta 2 TXTL. TXTL reactions were performed with eGFP templates (5 nM) for 8 hours at 30°C, and each fluorescence was measured at λ_ex_ 488 nm and λ_em_ 509 nm. (A) Akaby TXTL, (B) Akaby + Gams TXTL, (C) Rosetta 2 TXTL, and (D) Rosetta 2 + GamS TXTL. GamS (3.5 μM) was supplemented as a nuclease inhibitor. (E) The TXTL reaction comparison for eGFP fluorescence was generated with the T7Max-eGFP:BamHI template data. T7, T7 RNA polymerase promoter; T7Max, enhanced T7 RNA polymerase promoter; template named with BamHI, linearized plasmids by BamHI; NTC, no template control. The graphs show means with error bars that signify SEM (n=3). Significance was determined by Student’s t-Test. \*\**p* < 0.01.

With the eGFP gene under the regular T7 promoter, we were only able to detect a measurable fluorescence from plasmids. Neither Akaby nor Rosetta 2 TXTL generated a noticeable fluorescence from the linear eGFP templates with the T7 promoter (T7-eGFP:BamHI) (**Fig. 2A and C**). The linear eGFP templates with the T7Max promoter (T7Max-eGFP:BamHI) generated a fluorescent in Akaby TXTL (**Fig. 2A**), while the fluorescence observed in Rosetta 2 TXTL was as low as in the NTC (**Fig. 2C**).

Next, we used the established GamS addition protocol to help protect linear templates in TXTL[1]. We conducted experiments with the supplementation of GamS as a control in parallel (**Fig. 2B and D**). In Akaby TXTL, GamS supplementation did not produce a significant difference (**Fig. 2E**). On the contrary, in Rosetta 2 TXTL, we could see a significantly higher eGFP fluorescence from T7Max-eGFP:BamHI with the supplementation of GamS (**Fig. 2E**).

This demonstrated that Akaby TXTL improved protein expression from linear DNA templates, with the difference most apparent under a strong promoter.

### The abundance of mRNA from linear templates are increased in Akaby TXTL

The reduction of templates by nuclease activities may result in reducing its transcripts in TXTL. For further characterization of Akaby TXTL for protecting linear templates, we measured the amount of mRNA transcribed from eGFP templates and the eGFP fluorescence at several time points: 0, 1, 4, and 8 hours (**Fig. 3**).

**Fig.3.**
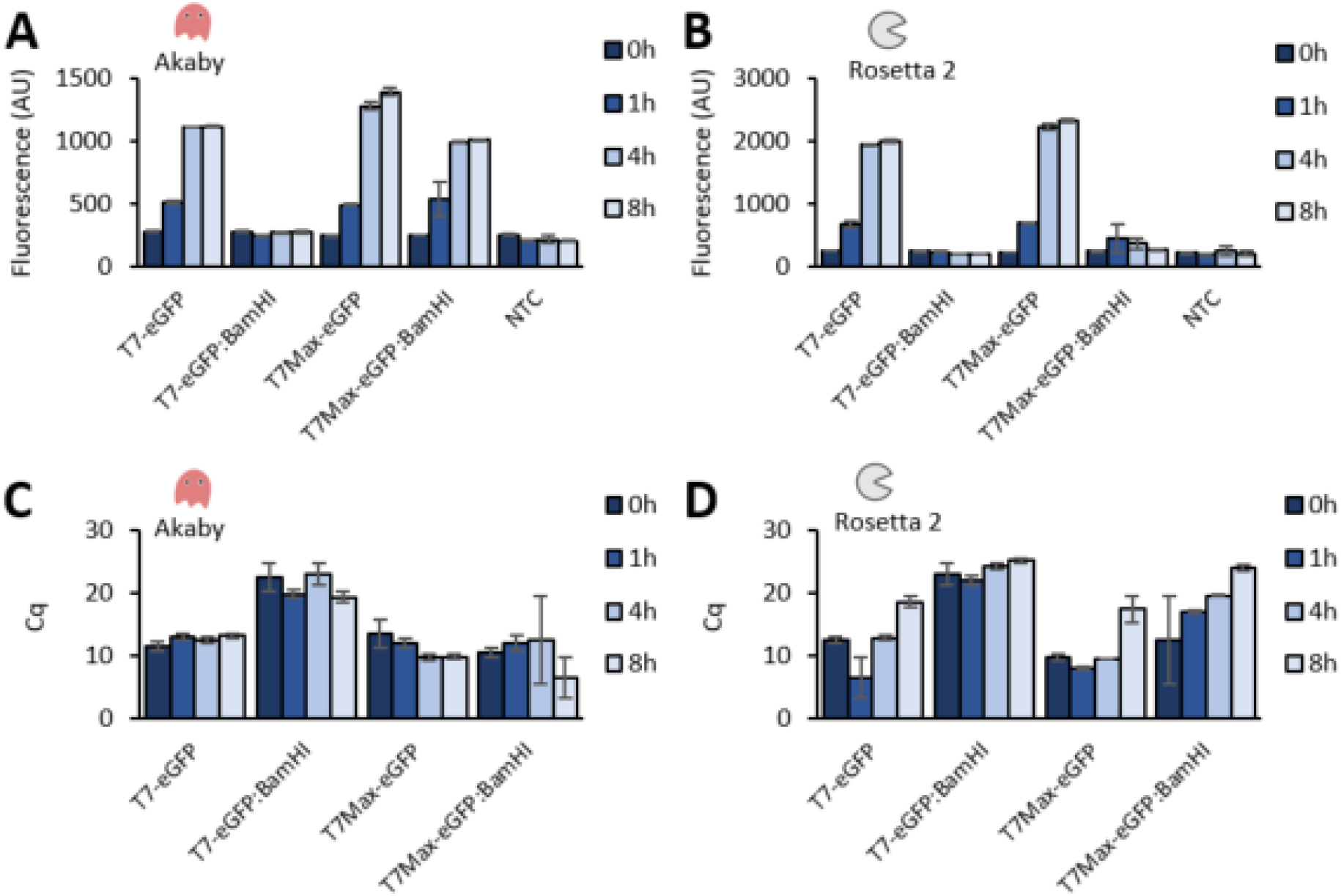
eGFP expression and mRNA abundance in Akaby and Rosetta 2 TXTL at 0, 1, 4, and 8 hours incubation. Fluorescence and mRNA abundance were measured at 0, 1, 4, and 8 hours. The TXTL reactions were incubated in a plate reader using black-bottom well plate. The measurements were performed with λ_ex_ 488 nm and λ_em_ 509 nm. (A) Fluorescence generated in Akaby TXTL and (B) in Rosetta 2 TXTL. Cq values of mRNA transcribed (C) in Akaby TXTL and (D) in Rosetta 2 TXTL. T7, T7 RNA polymerase promoter; T7Max, enhanced T7 RNA polymerase promoter; template named with BamHI, linearized plasmids by BamHI; NTC, no template control; Cq, quantitation cycle. The graphs show means with error bars that signify SEM (n=3).

The fluorescence generated from each template at 8 hours was consistent with the result we observed in **Fig. 2**. The eGFP production reached plateau around 4 hours of incubation (**Fig. 3A and B**). The quantitative reverse transcription PCR (RT-qPCR) established that mRNA abundance was consistent with the fluorescence measured from eGFP expression reactions (**Fig. 3C** and **D**).

In Akaby TXTL, all templates maintained a relatively high amount of mRNA over 8 hours, except T7-eGFP:BamHI (**Fig. 3C**). On the contrary, in Rosetta 2 TXTL, while the amounts of mRNA from plasmids were as high as in Akaby TXTL, the mRNA level from linear templates was almost the same as NTC, independent of the promoter sequences (**Fig. 3D**).

These data demonstrated that while Akaby TXTL increased linear template stability, the abundance of mRNA and the fluorescence also increased proportionally.

### Stability of short oligonucleotides without RBS in TXTL improves in Akaby

Since the initially investigated BamHI digested linear GFP DNA template was 3593 nt long, we decided to evaluate the effect of *RecB* knockout TXTL on shorter, untranslated DNA fragments. We designed 347 nt long DNA, which consisted of the truncated eGFP under the T7Max promoter, without the ribosome binding site. Therefore, the eGFP genes are not translated, but the DNA templates are still transcribed (**Fig. 4A**). This experiment was designed to eliminate the potential effect of the ribosome complex binding on mRNA stability.

**Fig.4.**
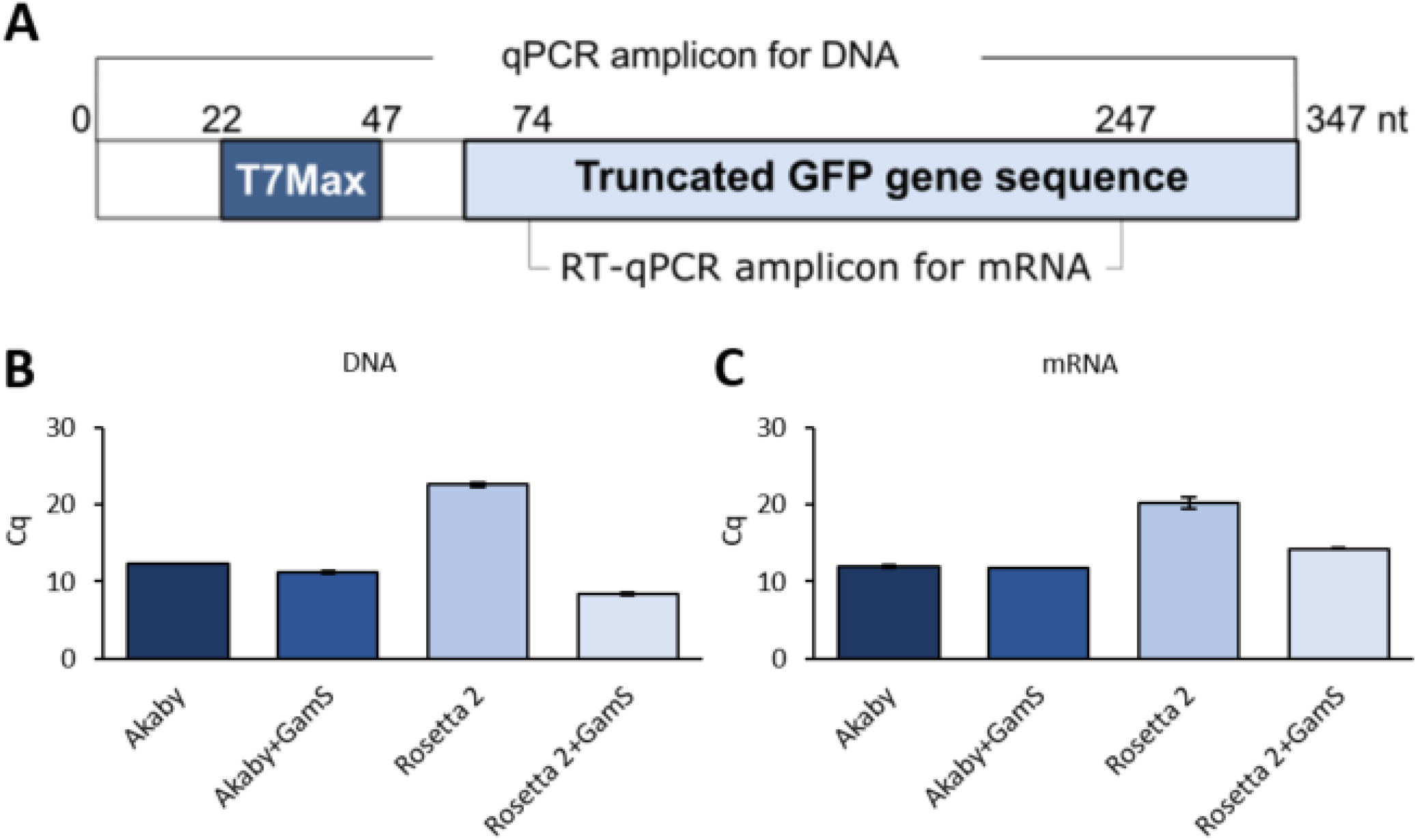
DNA and mRNA abundance from short DNA oligonucleotides in Akaby and Rosetta 2 TXTL at 8 hours incubation. (A) The design of a 347 nt length DNA oligo. The eGFP gene under the T7Max promoter is truncated by a stop codon insertion. This oligonucleotide does not contain the ribosome binding site. The amplicons sites for qPCR measurement are described: DNA (0-347 nt) and mRNA (74-247 nt). (B) The DNA oligo (5 nM) was incubated in Akaby or Rosetta 2 TXTL at 30°C for 8 hours. Then DNA was purified with a DNA miniprep kit and qPCR experiments were conducted. (C) The abundance of mRNA in TXTL after 8 hours incubation was measured by RT-qPCR. GamS (3.5 μM) was supplemented as a nuclease inhibitor. Cq: quantitation cycle. The graphs show means with error bars that signify SEM (n=3).

We incubated this short linear DNA in Akaby or Rosetta 2 TXTL for 4 and 8 hours, and measured the abundance of DNA and mRNA transcribed from the DNA fragments by qPCR and RT-qPCR accordingly (**SFig. 2** and **Fig. 4**).

After 8 hours of TXTL incubation, the abundance of the DNA fragment in Akaby TXTL was significantly higher than the one in Rosetta 2 TXTL, and GamS supplementation retained the highest DNA on both Akaby and Rosetta 2 TXTL (**Fig. 4B**). Consistent with the observation in DNA abundance, mRNA in Akaby TXTL was higher than in Rosetta 2 TXTL (**Fig. 4C**). Therefore, we concluded that Akaby TXTL protects linear DNA in the TXTL reaction, resulting in an increased transcription level of the templates.

## Discussion and Conclusions

### The efficiency of protein synthesis: Akaby vs Rosetta 2

Throughout the experiments, we observed the lower protein expression efficiency in Akaby cell-free transcription-translation (TXTL), compared to Rosetta 2 TXTL. TXTL is widely known to have batch variability in reaction rate, which can reason for this observation. Another explanation is that Rosetta 2 (a BL21 derivative) is a protease deficient strain designed for protein overexpression while Akaby (a DH5α derivative) is not. Since we initially had some experimental difficulties with BL21 strain, we shifted gears to using DH5α. We expect to see similar results with BL21 strains; however, since we already have a working strain for our TXTL reactions, we decided to leave out BL21 experimentation for this paper.

### Comparing abundance of eGFP templates

We compared the transcript abundance in Akaby TXTL under different conditions. At 8 hours, we saw mRNA abundance without GamS supplementation, except for T7-eGFP:BamHI (**Fig. 3**). However, GamS supplementation resulted in reducing the mRNA at 8 hours for all of the template types (**Fig. S1C**). Based on these RT-qPCR data, we speculate that GamS might negatively affect the sustainability of transcriptions in Akaby TXTL, however the exact mechanisms remains unknown. We saw eGFP fluorescence reach saturation around 4 hours, and thus we did not see differences with and without GamS supplementation in Akaby in fluorescence measurements (**Fig. 3A** and **SFig. 1A**).

In samples with the T7Max promoter, we observed that the similar transcript levels from both plasmids and linear templates in Akaby TXTL. However, the eGFP fluorescence was lower for the linear templates. We speculate this is due to nucleases that remain in Akaby TXTL (except for the deleted *RecB*)[24]. It is likely that the active nucleases in Akaby TXTL degrade the linear templates to some extent, reducing the full-length eGFP transcripts from the linear templates. In the RT-qPCR experiments, we only amplified the middle of the eGFP mRNA sequence. This may explain why we did not see clear differences in mRNA levels between the plasmids and the linear templates degraded on the edges (**Fig. 3**). Additionally, with the PCR amplified eGFP gene without long extensions on both ends, we could only see eGFP fluorescence with higher template concentration (100 nM) in Akaby TXTL (**SFig. 3** and **4**). This observation also supports the presence of other, less active nucleases.

In summary, we constructed a *RecB* knockout *E. coli* strain called Akaby, and we demonstrated a TXTL system based on this strain. Compared to standard TXTL Rosetta 2 extract, Akaby TXTL significantly improved linear DNA stability. Akaby matches the efficiency of GamS protection for linear DNA templates, without the need to purify additional proteins and add more components to the reaction mixture. Akaby TXTL can be used as a versatile TXTL platform for the expression of linear DNA templates.

In addition to the specific use case of linear DNA protection in Akaby, this work demonstrates proof of principle for a pipeline for producing *E. coli* TXTL strain mutants. As the use frequency and versatility of TXTL systems increases, this technique can be useful for engineering other *E. coli* strains for special TXTL applications.

## Acknowledgment

We thank Dr. Anna Corts for the helpful advice of homology recombineering in *E. coli*. We thank Dr. Richard Murray for sample of GamS protein. We thank Dr. Vincent Noireaux for helpful discussion and guidance in using the Chi6 system. This work was supported by NIH award 5R01MH114031, RNA Scaffolds for Cell Specific Multiplexed Neural Observation, NSF award 1840301, RoL:FELS:RAISE: Building and Modeling Synthetic Bacterial Cells, NSF award 1807461, SeMiSynBio Very Large scale genetic circuit design and automation program, and the Funai Foundation for Information Technology.

## Materials and methods

### *RecB* mutagenesis in *E. coli*

The protocol of Datsenko[21] with minor modifications was used for *RecB* disruption. Plasmids were purchased from Addgene. pKD4 (Addgene plasmid # 45605; http://n2t.net/addgene:45605; RRID:Addgene_45605) was used for Km^R^ template[21] and pKDsgRNA-trmI (Addgene plasmid # 89955; http://n2t.net/addgene:89955; RRID:Addgene_89955) was used for the lambda red recombinase system[25]. Primers were purchased from Integrated DNA Technologies, Inc. Km^R^ was PCR amplified with primers (HR FW and RV primer in Table. S1). These primers include 120-nt homology extensions that are complemental sequences of upstream or downstream of *RecB* in DH5α genome, and 19- or 21-nt priming sequences for pKD4 as a template. After PCR amplification of Km^R^, the reaction mixture was treated with DpnI, and then purified with *GenCatch* PCR Purification Kit (Epoch Life Science, Inc., No. 2360250). pKDsgRNA-trml was transformed into NEB® 5-alpha Competent *E. coli* (New England BioLabs Inc., C2987I). The 230 μl of pKDsgRNA-trml carrying *E. coli* pre-culture was inoculated in 35 ml SOB and incubated at 30°C. After 2 hours of incubation with OD_600_∼0.2, 150 μl 20% Arabinose was added to the culture to induce the lambda red recombinase system from the pKDsgRNA-trmI plasmid. When the OD_600_ reached 0.45, the cells were treated to be electrically competent. The amplified Km^R^ fragments flanking 120-nt homology arms were transformed into the cells. 1 ml SOC was immediately added after electroporation, and the culture was incubated at 37°C overnight. The following day, the entire culture was plated on LB plates containing 50 μg/ml Kanamycin, 200 μl culture per plate. The plates were incubated at 37°C overnight. The six *RecB* disrupted colonies were obtained. 10 μl total volume of colony PCR was performed with 0.4 μM of locus-(Primer 1 and 4) and insert-(Primer 2 and 3) specific primer pairs, 1X OneTaq® Quick-Load® 2X Master Mix with Standard Buffer (New England BioLabs Inc., M0486L), and 2 μl of water suspended colony. The colonies are stored as glycerol stocks, and one of them was used for cell extract preparation.

### Cell extract preparation and TXTL reaction

This protocol was adapted from Noireaux[23] and Jewett[26]. The Rosetta 2 cell extract preparation was followed by the method described previously[27] with one modification. A 750 ml 2xYPTG was grown at 30°C instead of 37°C. Akaby extract preparation was performed with the same protocol of Rosetta 2 cell extract preparation with minor modifications. For the starter culture preparation, a 50 ml starter culture of the Akaby was grown to saturation at 30°C in 2xYPTG with Kanamycin (50 μg/ml). A 750 ml of 2xYPTG culture (without antibiotic) was inoculated with a 10 ml starter culture. The culture was grown at 30°C to an OD_600_ of 0.4-0.6, then harvested.

Cell-free transcription-translation (TXTL) reaction was performed based on the method described previously[27]. Unless otherwise specified, the template concentration was 5 nM. GamS was added with a concentration of 3.5 μM. The TXTL reaction was incubated at 30°C for 8 hours, followed by 4°C temperature hold. For linear eGFP templates preparations, eGFP plasmids were digested with BamHI (New England BioLabs Inc., R0136L) at 37°C for 1 hour and agarose gel-purified with *GenCatch* Advanced Gel Extraction Kit (Epoch Life Science, Inc., No. 2260050). eGFP fluorescence was measured at λ_ex_ 488 nm and λ_em_ 509 nm with plate reader PMT setting “medium” and 6 reads per well. All fluorescent measurements were performed on SpectraMax. For the endpoint measurement, 14 μl of TXTL reaction was transported into a 384 black bottom well plate to measure after the incubation. For kinetics, 15 μl of TXTL was prepared in a 384 black bottom well plate, sealed with clear sealing tape to avoid evaporation, and measured every hour, including at the start of the incubation of 30°C.

### Relative comparison of transcripts with Reverse Transcription-quantitative Polymerase Chain Reaction (RT-qPCR)

Template DNA in 2 μl of the TXTL reaction was degraded by adding 0.5 μl of TURBO DNase (2 U/μl, Catalog No. AM2238, Invitrogen). The mixture was incubated at 37°C for 30 minutes. The enzyme and the expressed proteins were inactivated by adding 15 mM EDTA (Catalog No. E9884, Sigma-Aldrich) at 75°C for 10 minutes (T100 Thermal Cycler, Bio-Rad). The denatured proteins were pelleted through centrifugation at 3,200 *g* for 2 minutes. For DNA abundance measurement in TXTL, DNA in 20 μl of TXTL reaction was purified with *GenCatch* PCR Purification Kit before reverse transcription, instead of DNase treatment. The DNA fragment binding to the column membrane was eluted with 20 μl water. The elution was re-applied to the column membrane and repeated the elution 2 more times. A 2 μl of the final elution was used for the reverse transcription reaction.

For each protein sample, forward and reverse primers (Integrated DNA Technologies) were created for downstream reverse transcription and qPCR experiments. Each primer pair was compatible with transcripts produced from the regular promoter and T7Max and wild-type cell extract and Akaby cell extract. For eGFP, the forward primer was Primer 5 and the reverse primer was Primer 6. For the short eGFP transcript that was fragmented due to an integrated stop codon in the gene, the forward primer was Primer 7 and the reverse primer was Primer 8.

To prepare the reverse transcription reaction, 2 μl of the DNase-treated sample was mixed with 2 μl of 10 μM reverse primer, 4 μl of 5X Protoscript II Reverse Transcriptase Buffer, 1 μl of Protoscript II Reverse Transcriptase (200 U/μl, Catalog No. M0368, New England BioLabs Inc.), 2 μl of 0.1M dithiothreitol (DTT), 1 μl of 10mM dNTP, 0.2 μl of RNase Inhibitor (Catalog No. M0314, New England BioLabs Inc.), and 8 μl of nuclease-free water. The reverse transcription reaction was incubated at 42°C for 1 hour and the reverse transcriptase was inactivated at 65°C for 20 minutes.

The quantitative PCR reaction mix was prepared by mixing 2 μl of complementary DNA from the reverse transcription with 2 μl of 10 μM forward and reverse primers, 11.25 μl OneTaq Hot Start 2X Master Mix with Standard Buffer (Catalog No. M0484, New England BioLabs Inc.), 1.25 μl Chai Green Dye 20X (Catalog No. R01200, Chai Bio), and 7.5 μl of nuclease-free water. The RT-qPCR was completed using Open qPCR (Chai Biotechnologies) with the following thermocycling program: 1 cycle of 30 second denaturation at 95°C, 30 cycles of 15 second denaturation at 95°C, 15 second annealing at 50°C, 1 minute extension at 68°C, and 1 cycle of 5 minutes final extension at 68°C. The amplification curves plotted through the Open qPCR software to determine Cq values and averages across 3 replicates of each promoter type were calculated separately.

### Short DNA fragment preparation

The oligo template and primers (Primer 9 and Primer 10) for the amplification (Sequence is in STable 1) were purchased from Integrated DNA Technologies, Inc. The oligo was PCR amplified with OneTaq® 2X Master Mix with Standard Buffer (New England BioLabs Inc., M0482L), followed by agarose gel purification with *GenCatch* Advanced Gel Extraction Kit (Epoch Life Science, Inc., No. 2260050).

## Abbreviations

TXTL: cell-free transcription-translation
RT-qPCR: quantitative reverse transcription PCR

## Supplemental Information

**(SFigure 1).**
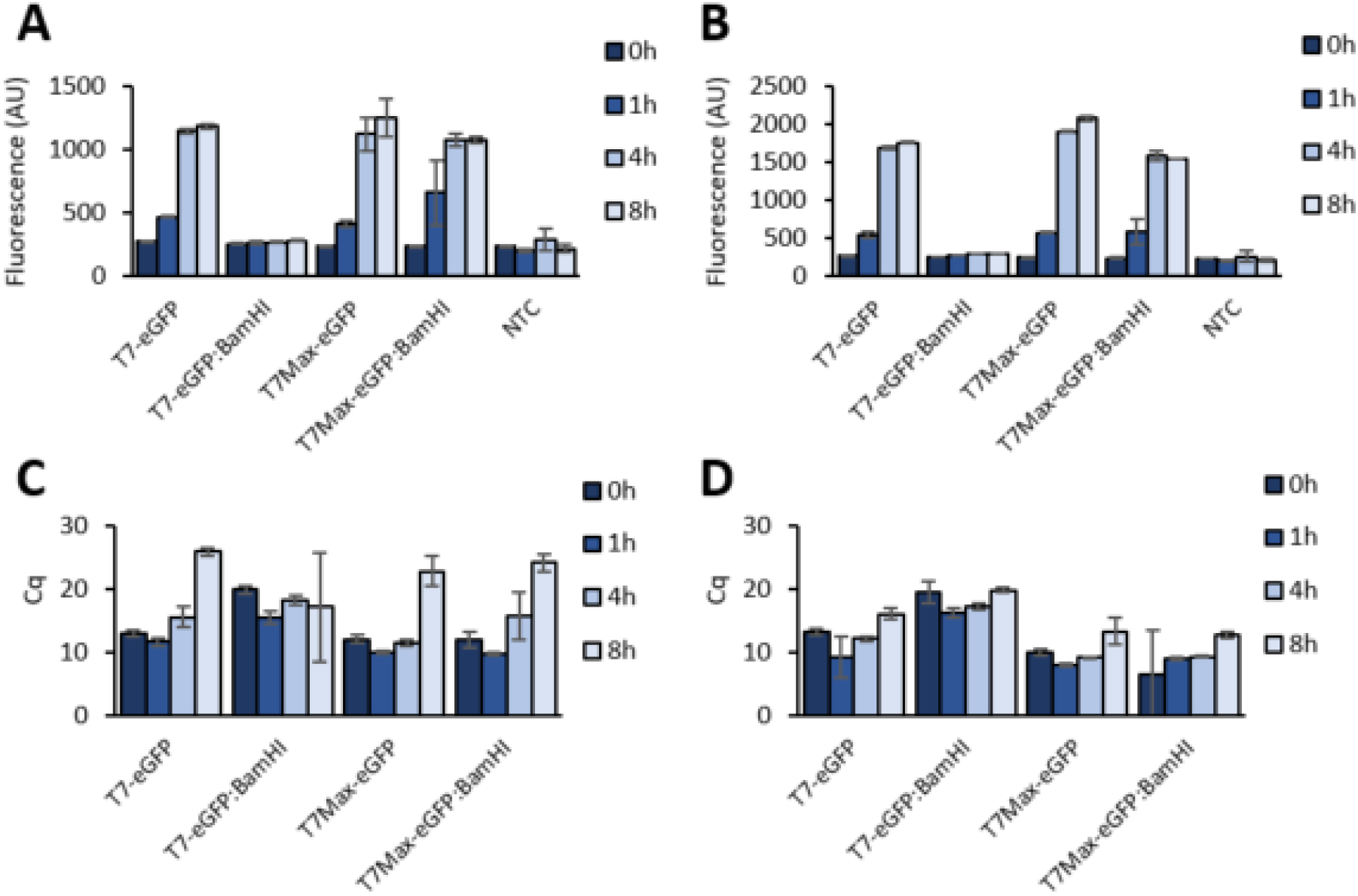
eGFP expression and mRNA abundance in Akaby and Rosetta 2 TXTL with the supplementation of GamS at 0, 1, 4, and 8 hours incubation. Fluorescence and mRNA abundance were measured at 0, 1, 4, and 8 hours. The TXTL reactions were incubated in a black-bottom well plate in a plate reader measuring at λ_ex_ 488 nm and λ_em_ 509 nm. (A) Fluorescence generated in Akaby + GamS TXTL and (B) in Rosetta 2 + GamS TXTL. For the mRNA measurement, TXTL reactions are incubated in a thermal cycler. At each time point, 2 μl of TXTL was taken, and RT-qPCR was performed. (C) The Cq values of mRNA transcribed in Akaby + GamS TXTL and (B) in Rosetta 2 + GamS TXTL. T7, T7 RNA polymerase promoter; T7Max, enhanced T7 RNA polymerase promoter; template name with BamHI, lineariws plasmids by BamHI; Cq, quantitation cycle. The graphs show means with error bars that signify SEM (n=3).

**(SFigure 2).**
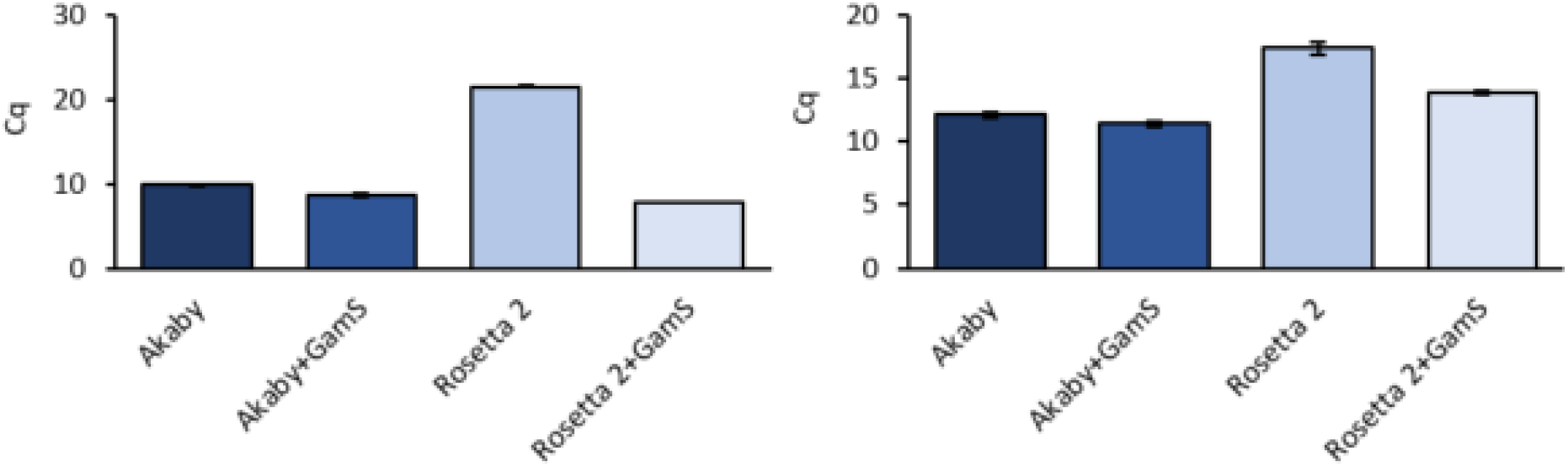
DNA and mRNA abundance from short DNA oligonucleotides in Akaby and Rosetta 2 TXTL at 4 hours incubation. (Left) The DNA oligo (5 nM) was incubated in Akaby or Rosetta 2 TXTL at 30°C for 4 hours. The DNA in TXTL reactions was purified with DNA miniprep kit, then qPCR was conducted. (Right) The abundance of mRNA in each TXTL reaction after 4 hours of incubation was measured by RT-qPCR. GamS (3.5 μM) was supplemented as a nuclease inhibitor. Cq: quantitation cycle. The graphs show means with error bars that signify SEM (n=3).

**(SFigure 3).**
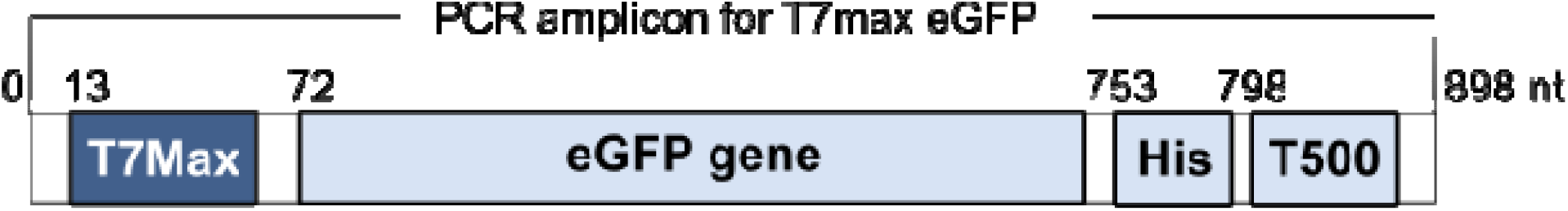
PCR amplified eGFP gene construct. The design of a 898 nt length DNA oligo. The oligo was PCR amplified from T7Max-GFP plasmid with Primer 11 and Primer 12 with LongAmp Taq 2X Master Mix (New England BioLabs Inc., M0287S). T7Max, the enhanced T7 polymerase promoter sequence; eGFP gene, an eGFP coding sequence; His, a 8xHistidine taq sequence; T500, a transcription terminator sequence.

**(SFigure 4).**
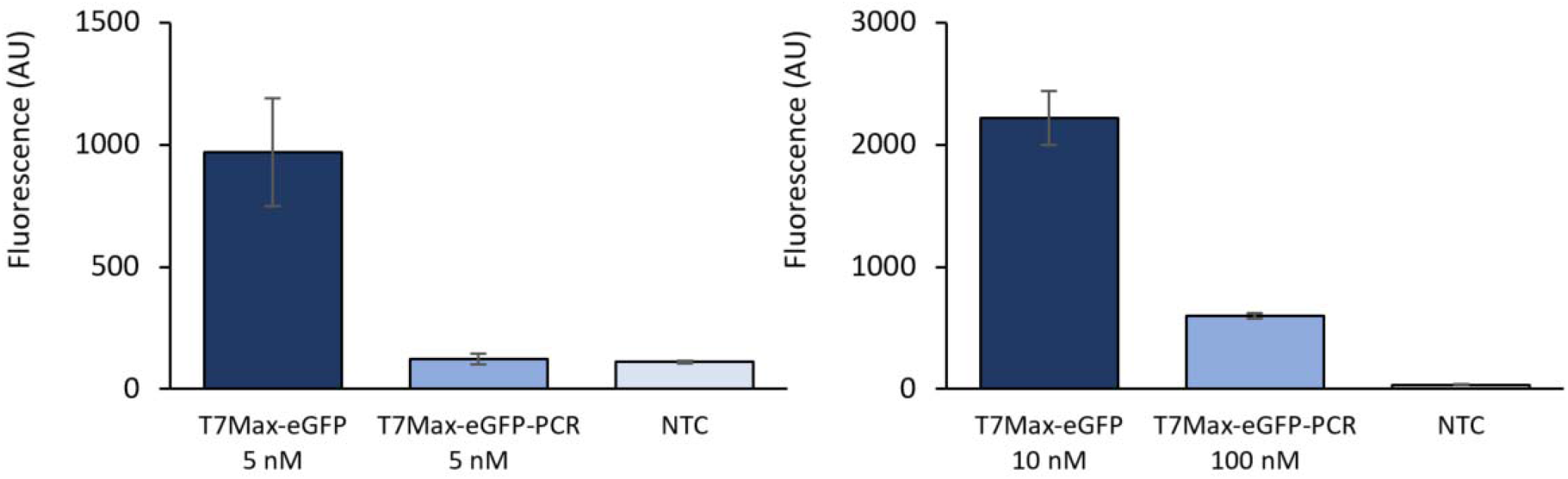
eGFP expression from PCR amplified linear eGFP gene templates. The eGFP fluorescence was measured at λ_ex_ 488 nm and λ_em_ 509 nm after 8 hours of incubation. The TXTL reaction was performed in Akaby TXTL (A) with 5 nM PCR amplified templates and (B) with 10 nM plasmid concentration and 100 nM PCR amplified template. T7Max-eGFP, eGFP coding plasmids containing T7Max promoter; T7Max-eGFP-PCR, PCR amplified eGFP gene fragments containing T7Max promoter; NTC, no template control. The graphs show means with error bars that signify SEM (n=3).

**(STable 1).**
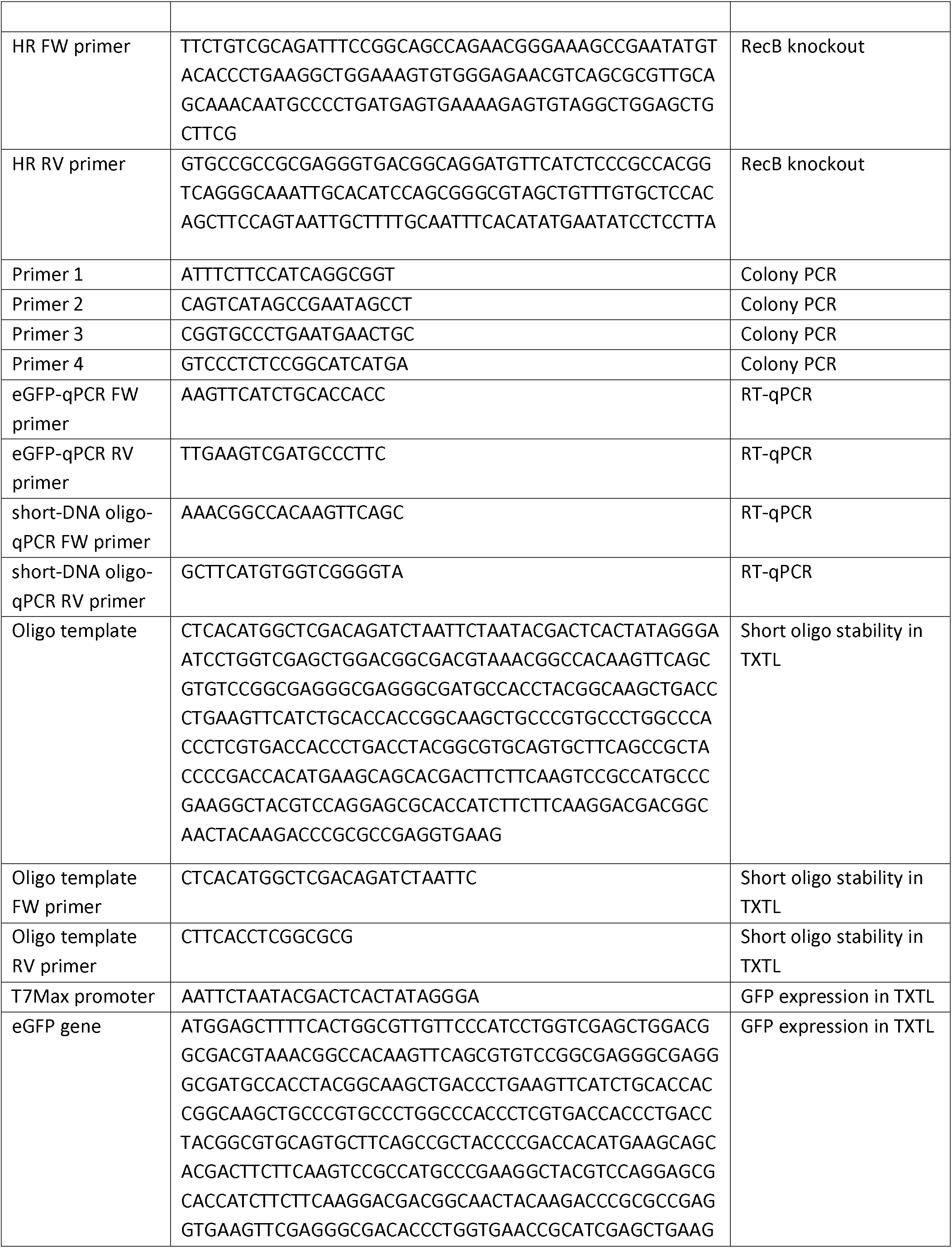
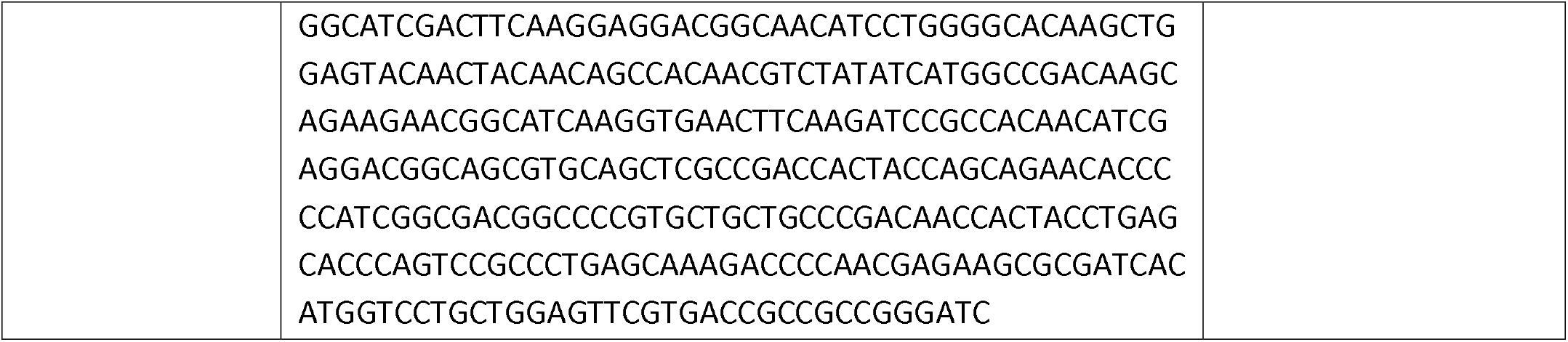
Oligonucleotide sequences.

**(STable 2).** Plasmid information.

| Name | Purpose |
| --- | --- |
| pKDsgRNA-trml | RecB knockout |
| pKD4 | RecB knockout |
| T7-GFP | GFP expression in TXTL |
| T7Max-GFP | GFP expression in TXTL |

